# Fast Virome Explorer: A Pipeline for Virus and Phage Identification and Abundance Profiling in Metagenomics Data

**DOI:** 10.1101/196998

**Authors:** Saima Sultana Tithi, Roderick V. Jensen, Liqing Zhang

## Abstract

Identifying viruses and phages in a metagenomics sample has important implication in improving human health, preventing viral outbreaks, and developing personalized medicine. With the rapid increase in data files generated by next generation sequencing, existing tools for identifying and annotating viruses and phages in metagenomics samples suffer from expensive running time. In this paper, we developed a stand-alone pipeline, FastViromeExplorer, for rapid identification and abundance quantification of viruses and phages in big metagenomic data. Both real and simulated data validated FastViromeExplorer as a reliable tool to accurately identify viruses and their abundances in large data, as well as in a time efficient manner.

## 1 Introduction

Identifying the kinds of viruses that infect eukaryotes and prokaryotes (phages) and understanding their functions are important because they are the most abundant entities on Earth [18]. Even in a healthy human body, there are estimated to be 100 times more viral particles than eukaryotic cells [7]. Studies have shown that there are connections between human gut microbiome (viruses and bacteria) and diseases such as diabetes [6] and depression [8] and cancer [31]. Moreover, recent emerging viral outbreaks including Zika outbreak in Brazil [3], Ebola in West Africa [4, 9], Middle East respiratory syndrome coronavirus (MERS-CoV) [10], SARS, influenza-A caused tens of thousands of human deaths. To better understand and eventually prevent such viral outbreaks, it is critical to have timely identification and annotation of viruses. Traditional techniques of virus identification rely on isolation and clone culturing, which is not only time-consuming but often infeasible as many viruses and their hosts are difficult to cultivate in laboratories. Thanks to the fast development of biotechnology, it is now easy and quick to produce metagenomics data for a direct analysis of genetic materials to identify viruses and their abundances in various environments [11].

However, with the ease of metagenomics data generation also comes the challenge of downstream data analysis, including the computational identification of viral species and their abundances in a fast yet accurate manner from hundreds of millions/billions of short sequences. Strategies to identify and annotate viruses vary among different tools, ranging from analyzing marker genes, binning sequences or reads into taxonomic groups, assembling sequences into contigs and then annotating the taxonomy using the contigs, to directly aligning short reads to a reference database and inferring virus types and abundances based on the alignment results. The most straightforward and fastest approach for virus taxonomic annotation is to align short reads to a marker gene database and identify viruses based on the alignments, for example, MetaPhlAn [26] and its updated version MetaPhlAn2 [29] use this approach. However, marker gene analysis strategy does not work well when the input data contain species that do not have known marker genes. Comparatively, assembling reads into longer contigs and then performing the taxonomic analysis with contigs tend to produce more accurate results [21]. This type of virus analysis pipelines normally requires the users to assemble the reads using an independent assembler and then annotates the assembled contigs (e.g., VirSorter [22], Metavir [23], Metavir2 [24], and Virome [30]). Understandably the assembly of short reads into contigs gives longer sequences including longer coding regions with more informative content, which leads to improved annotation and downstream analysis. However, read assembly can be very time-consuming for large metagenomics data and can also generate chimeras (i.e., sequences from different genomes that are incorrectly assembled together due to their similarity) that mislead downstream annotation. Finally, tools such as MG-RAST [16], ViromeScan [20], VIP [15], and HoloVir [12] directly align short reads to a reference database of whole genomes for taxonomy annotation. Many of these tools were initially developed for bacteria but adapted later for viruses and tend to work poorly due to the much smaller reference databases available for viruses than for bacteria [7]. In addition, as many virus annotation tools (i.e., Metavir [23], Metavir2 [24], Virome [30], MG-RAST [16]) are web-based, users need to upload their data to the website and wait for a long time to get results.

To provide fast and accurate virus annotation on metagenomics data, we developed a stand-alone pipeline, FastViromeExplorer. Instead of the traditional read alignment tools such as BLAST [1] or Bowtie2 [13], FastViromeExplorer uses kallisto [2], a pseudoalignment based approach originally developed for alignment and quantification of RNA-seq data, to rapidly map short reads to a reference virus database. Then it filters the alignment results and reports virus types and abundances along with taxonomic annotation. To test the performance of FastViromeExplorer, we used simulated datasets of a known mixture of viral, phage, and bacterial genomes with different error/mutation rates. We also applied FastViromeExplorer to real metagenome datasets generated from a Fecal Microbiota Transplantation (FMT) experiment [14]. FastViromeExplorer is directly compared with the gold standard tool Blastn, and with ViromeScan [20], a recently developed read based annotation tool for eukaryotic viruses.

FastViromeExplorer is freely available at https://code.vt.edu/saima5/FastViromeExplorer.

## 2 Methods

FastViromeExplorer, written in Java, has two main steps, (1) the read mapping step where all reads are mapped to a reference database, and (2) the filtering step where the mapping results are subjected to three major filters (detailed later) for output of the final annotation results on virus types and abundances. The input of the read alignment step is raw reads (single-end or paired-end) in fastq format. FastViromeExplorer uses the reference database downloaded from NCBI containing 8,957 RefSeq viral genomes as default but can also use any updated or customized databases as reference. A pre-computed kallisto index file, generated for the 8,957 genomes, is distributed with FastViromeExplorer.

First, FastViromeExplorer calls kallisto [2] as a subprocess to map the input reads against the reference database. Kallisto was developed to map RNA-seq data to a reference transcriptome (all the transcripts for a genome) leveraging the pseudoalignment process and estimate the abundance of the transcripts using the Expectation-Maximization (EM) algorithm [5]. As there is no actual sequence alignment of the entire read over the reference sequences, the pseudoalignment process enables read mapping to be both lightweight and superfast. For example, kallisto was able to map and quantify 30 million paired-end RNA-seq reads for the human transcriptome in less than 10 minutes on a small laptop computer with a 1.3-GHz processor [2]. In addition to the ultrafast speed, kallisto also gives accurate estimation of abundance of each transcript or reference sequence [25, 28]. Consequently, kallisto could provide an ideal tool for virus type and abundance annotation in metagenomics samples that commonly have tens of millions of reads, mapping of which using commonly used programs such as BLAST can be time-consuming and often infeasible without computer clusters. Therefore, FastViromeExplorer deploys kallisto for the purpose of read mapping and abundance estimation of the annotated viruses. The k-mer size in kallisto can be altered depending on user’s need, and experiments have shown that the default size 31 works well for metagenomics data and is therefore kept as default also in FastViromeExplorer.

After the first alignment step, FastViromeExplorer takes the output of kallisto that includes information of the aligned reads together with estimated abundances or estimated read counts of all the identified viruses for the processing of the second step. The second step filters the output of the first step using three criteria, introduced to ensure the annotation quality and especially to reduce the number of false positive annotations. In detail, the first criterion, hereafter referred to as “*R*”, is based on the ratio of the observed extent of genome coverage with the expected extent of genome coverage, computed as

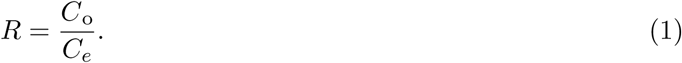

*C*_*o*_ is the observed extent of genome coverage by the mapped reads, computed as

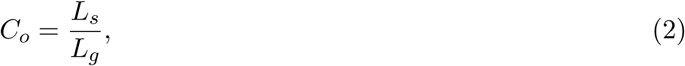

where *L*_*s*_ is the actual length of the genome that is supported or covered by the mapped reads and *L*_*g*_ is the length of the genome. *C*_*e*_ is the expected extent of genome coverage, assuming a Poisson distribution for the mapped reads along the genome, and therefore,

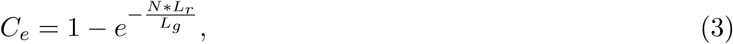

where *N* is the number of mapped reads to the genome, *L*_*r*_ is the read length, and *L*_*g*_ is the length of the genome. If a virus has *R <* 0.3, FastViromeExplorer discards the virus. This criterion is motivated by the observation that some annotated viruses only have reads mapped to the repeat regions of their genomes. For example, while analyzing the fecal samples from Lee et al. [14], we found that for the BeAn 58058 virus (NC 032111.1), all the reads were mapped to one particular region of its genome, from 8,200 bp to 8,700 bp (see supplementary figure 1). Analyzing this region using RepeatMasker [27] revealed that it is a simple repeat region and falls into the class of Alu elements. If the virus is truly present in the sample, we expect reads to be mapped to not only the repeat region but also other regions of the genome. Therefore, finding this virus is likely an artifact caused by the prevalence of repeat regions instead of real biological signals. By imposing this criterion, we filter out any virus for which all the reads are mapped to a repeat region of the virus genome. If a virus is truly present in the sample, the mapped reads to the virus should come from random locations of the genome, assuming that the mapped reads follow a Poisson distribution along the genome, the expected coverage of the virus genome *C*_*e*_ can be computed by Equation (3). If the reads are all mapped to a repeat region, the observed coverage of the virus genome *C*_*o*_ is expected to be much lower than *C*_*e*_, as a result, *R* is low and by imposing a cutoff of 0.3 (determined based on our empirical analyses), viruses that have reads mapped to only repeat regions get filtered out.

**Figure 1:**
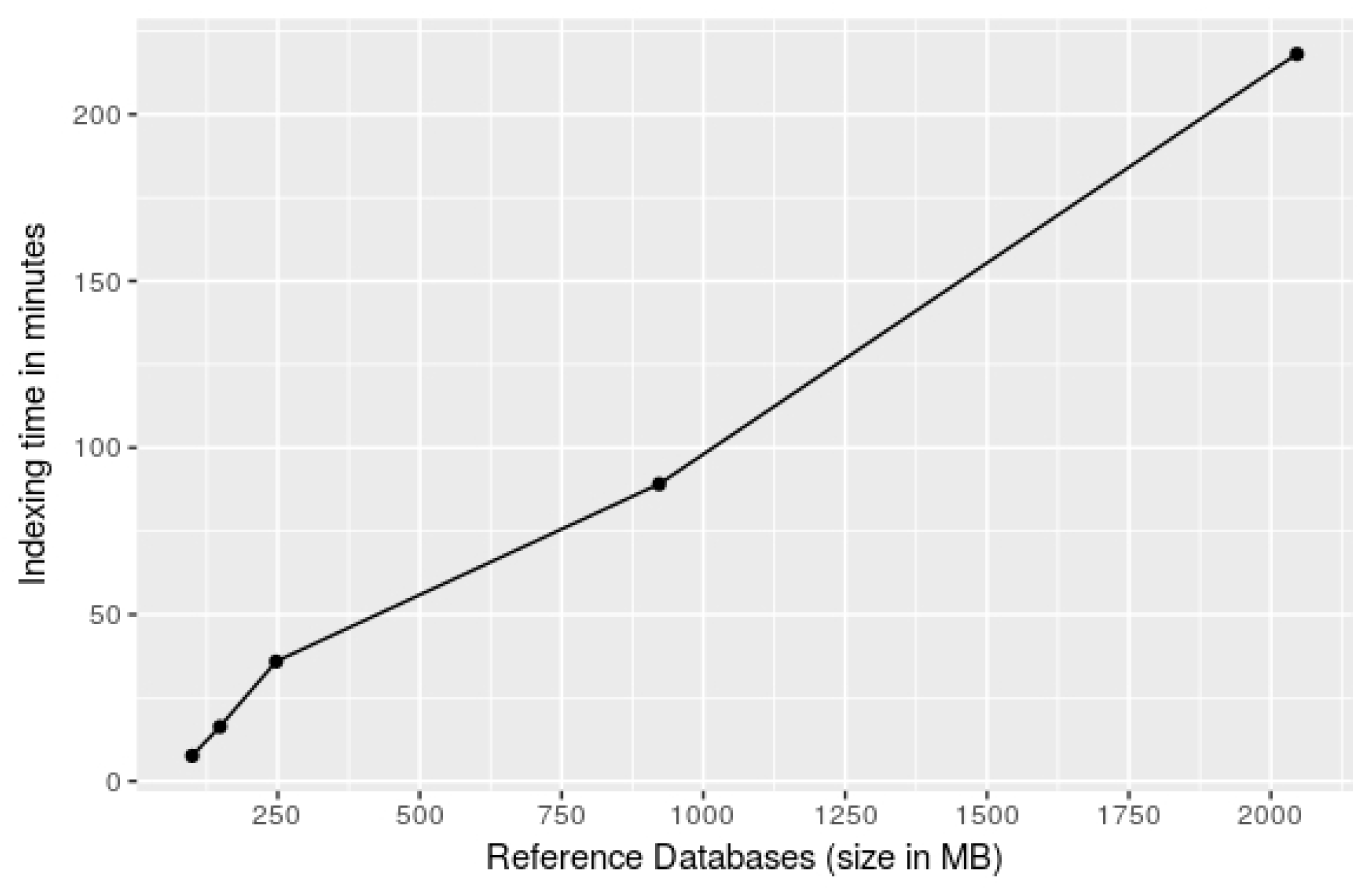
Kallisto’s indexing time for five reference databases, NCBI RefSeq Eukaryotic viruses (99 MB), NCBI RefSeq Phages (148 MB), All NCBI RefSeq viruses and phages (247 MB), 62,921 mVCs (992 MB), and 125,842 mVCs (2 GB).

The second criterion requires *C*_*o*_ *≥* 0.1, that is, a virus that has *C*_*o*_ *<* 0.1 is discarded. This criterion requires that the mapped reads should cover at least 10% of the viral genome. Manual inspection of the annotation results reveals that very large viruses may have several repeat regions in their genomes and as a result, though all the reads are mapped to the repeat regions, they are mapped to different repeat regions. In these cases, the difference between *C*_*o*_ and *C*_*e*_ may be small and therefore *R* can be high enough to pass the first filter. However, it is very likely that the annotation is simply an artifact of repetitive sequences. For example, while analyzing the fecal samples [14], we found that Pandoravirus dulcis (NC 021858.1), a very large virus with 1,908,524 bps, has several repeat regions, and all the reads were mapped only to the repeat regions (see supplementary figure 2). Hence, to alleviate this annotation artifact, *C*_*o*_ *≥* 0.1 is used as the second filter. As repeat regions of a virus usually cover less than 10% of the genome [19], if any virus is covered for more than 10% by the reads, it is reasonable to assume that the reads are not merely from repeat regions and thus the virus should be considered in the annotation result.

**Figure 2:**
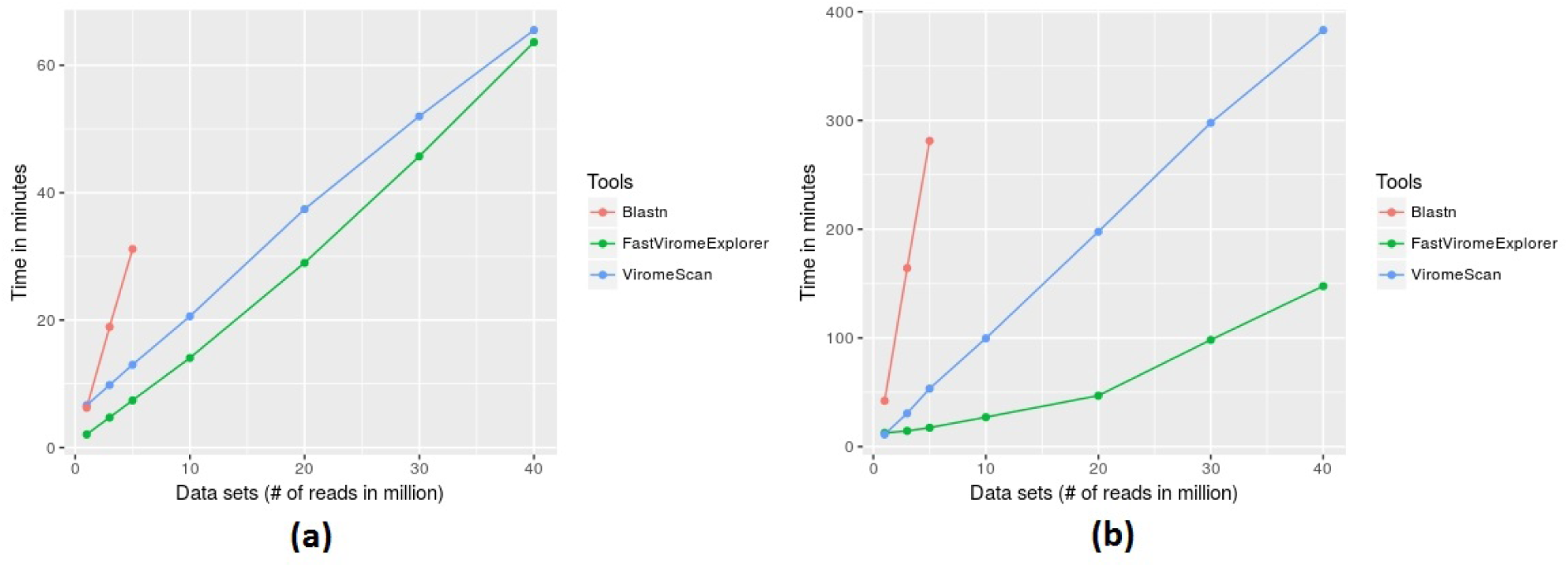
Comparison of running time among FastViromeExplorer, ViromeScan, and Blastn for seven data sets with 1, 3, 5, 10, 20, 30, and 40 million reads, respectively (a) against a reference database containing 8,957 NCBI RefSeq viruses, (b) against a reference database containing 125,842 mVCs

The third criterion is based on the number of mapped reads *N*. Extensive empirical analysis and inspection of the annotation results show that for very small viruses, only a few reads are enough to cover a good portion of the viral genome, resulting in high *R* and *C*_*o*_ that pass criteria 1 and 2. For example, in the fecal samples [14] that we annotated, four reads were mapped to Rose rosette virus RNA3 (NC_015300.1). As the virus has only 1,544 bps, four reads of length 150 bps were enough to pass criteria 1 and 2. But as only a handful of reads are mapped, it is likely that the virus is false positive. To be more stringent, FastViromeExplorer applies the third filter requiring the number of mapped reads to be greater than 10, and therefore discards the ones with *N <* 10.

After applying all the filters, FastViromeExplorer outputs the final result that contains a list of identified viruses in the given sample along with the estimated read count or abundance and taxonomy of the viruses. The output list is sorted by the abundance with the most abundant viruses on the top of the list.

It is worth noting that the three criteria are introduced to alleviate annotation artifacts caused by factors such as repeat sequences and low genome coverage, the actual cutoff values for *R*, *C*_*o*_, and *N* are based on our empirical experience and literature observation, and depending on the specific studies and the need of users, the cutoff values used here might not be suitable. To allow for flexibility and customization, FastViromeExplorer incorporates these three filters as parameters so that users can easily adjust the values to adapt to their own studies. For example, users can deploy more stringent criteria by setting higher values for *R*, *C*_*o*_, and *N* than the default, to get a “high confidence” set of viruses.

FastViromeExplorer was run on both simulated and real data to examine its running time and accuracy. FastViromeExplorer used kallisto (version 0.43.1) with default settings and generated pseudoalignment results in sam format and filtered abundance results in a tab-delimited file. The abundance results contain identified virus names, NCBI accession numbers, NCBI taxonomic path, and estimated read counts. FastViromeExplorer was run on two different reference databases, the default database distributed together with FastViromeExplorer, that is, the NCBI RefSeq database containing 8,957 genomes of eukaryotic viruses and phages, and the set of sequences collected from the JGI “earth virome” study [18] containing 125,842 metagenomic viral contigs (mVCs). The taxonomic annotation and host information for these mVCs were collected from the IMG/VR database [17].

In addition to the challenge of mapping 10s or 100s of millions of metagenomic reads, tools for the accurate identification and quantification of viral genomes must also be capable of handling ever-growing reference databases of viral sequences. In order to measure how the indexing step of kallisto scales with reference databases of different sizes, kallisto was applied to index five different databases. Three databases were generated from NCBI RefSeq viral database, one containing only phages (2,187 phage genomes), one containing only eukaryotic viruses (6,770 eukaryotic virus genomes), and one containing both phages and eukaryotic viruses (8,957 viral genomes). The other two databases were created from sequences collected from Paez-Espino et al. [18], one containing all the 125,842 mVCs and the other containing half of the mVCs. The time analysis of kallisto’s indexing step was produced on a Linux based cluster with 64 CPUs and 128 GB RAM. The indexing step was run using default k-mer size 31 and default number of threads 1. The precomputed kallisto index file for the full 125,842 mVCs from JGI is available here: https://bioinformatics.cs.vt.edu/zhanglab/software/FastViromeExplorer/.

To evaluate the performance of FastViromeExplorer, we compared speed and accuracy with ViromeScan, a recently developed virus annotation pipeline that calls BOWTIE2 as a subprocess for read mapping, that was shown to be 1,000 times faster than previous tools [20]. ViromeScan was run with default settings and with the eukaryotic DNA/RNA virus database containing 4,370 genome sequences, the largest reference database provided by ViromeScan, and with a custom database consisting of the 125,842 mVCs from JGI. ViromeScan generated alignment results and abundances of viruses at family, genus, and species level. We also ran Blastn (version ncbi-blast-2.6.0 +) using both the NCBI RefSeq viral database and the large JGI database. Blastn only generated the alignment result in text format. All the time analyses were calculated using elapsed real time from Unix’s time command.

To examine the annotation accuracy of FastViromeExplorer, simulated metagenomic data were used. A randomly selected collection of genomes containing 4000 virus genomes and 2000 bacteria genomes were obtained from NCBI RefSeq database. Four paired-end read datasets, each containing one million reads of length 100bps, were generated from these genomes using the read simulator WGSIM (https://github.com/lh3/wgsim). For all the datasets, 49% reads were from viruses and 51% from bacteria. The four datasets were generated using 1% sequencing error rate and 3%, 5%, 7%, or 10% mutation rates respectively. ViromeScan and Blastn were also applied to these four datasets. As ViromeScan uses eukaryotic viruses as the reference database, for comparison, both FastViromeExplorer and Blastn were run on a reference database containing only NCBI RefSeq eukaryotic viruses. ViromeScan was run with the eukaryotic virus database provided by ViromeScan. Under the default setting, ViromeScan removed all the mapped reads during its quality filtering and trimming step (trimBWAstyle.pl script) and did not produce any results. Therefore, it was run without ViromeScan’s quality filtering and trimming step. With the ground truth for the alignment of the reads, recall, precision, and F1 score were calculated using the following formula:

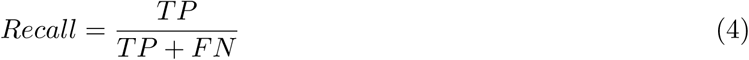

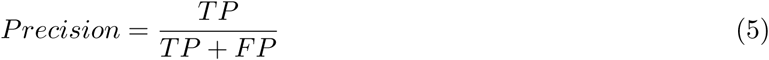

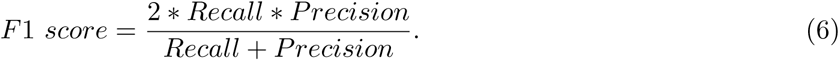

To examine the running time and annotation performance of FastViromeExplorer on real data, the fecal metagenomics datasets described in Lee et al. [14] were downloaded from NCBI under the accession number SRP093449 and annotated with both FastViromeExplorer and ViromeScan. The study tracked bacteria colonization in a fecal microbiota transplantation (FMT) experiment through analyzing metagenomics data. To examine how the viruses/bacteriaphages were affected by the transplantation, we reanalyzed the four fecal metagenomic samples collected from a healthy donor and three samples from a recipient patient suffering mild/moderate ulcerative colitis. The three samples for the recipient were collected prior to FMT, four weeks after FMT, and eight weeks after FMT, respectively. All the reads were Illumina paired-end reads with 150bp read length. Seven data sets of different sizes (1, 3, 5, 10, 20, 30, and 40 million reads) were also generated from the samples and annotated by FastViromeExplorer and ViromeScan to compare their running time on large datasets. To examine the effect of the reference database on annotation results, FastViromeExplorer was applied to the samples using two different reference databases, FastViromeExplorer’s default reference database and the set of 125,842 mVCs collected from the study [18]. While using the NCBI RefSeq database as reference, a Linux based laptop with Intel core i5-3230M CPU @ 2.60 GHz * 4 processors and 12 GB RAM was used to produce the results, and while using the 125,842 mVCs as reference, a Linux based cluster with 64 CPUs and 128 GB RAM was used to produce the results. While using the cluster, only 1 thread was used to run the tools.

## 3 Results and Discussion

We applied kallisto to index five databases of different sizes and calculated the running time of the indexing step. Figure 1 shows that indexing time increases linearly with the size of the reference databases, and for the largest reference database of 2 GB, kallisto took 3 hours and 38 minutes to generate the index file.

To examine how running time changes with sample size, we created seven data sets with 1, 3, 5, 10, 20, 30, and 40 million reads respectively from the data described in Lee et al. [14] and applied FastViromeExplorer, ViromeScan, and Blastn. As Blastn took too long to run on large data sets, we run Blastn on only three data sets of size 1, 3, and 5 million reads respectively. Two databases, one containing all NCBI RefSeq viral genomes and the other containing 125,842 mVCs from Paez-Espino et al. [18], were used as the reference databases, to also examine the effect of reference databases on running time. Figure 2a shows the running time using the NCBI database as reference. FastViromeExplorer has the shortest running time for all the seven data sets. For the data set with 5 million reads FastViromeExplorer took only seven minutes, compared to 12 minutes for ViromeScan, 31 minutes for Blastn. The speedup of FastViromeExplorer compared to ViromeScan became much more pronouced when a larger reference database was used. Figure 2b shows that when we used the larger reference database, for a data set with 5 million reads, FastViromeExplorer took 17 minutes, compared to 53 minutes for ViromeScan, and 4 hours and 40 minutes for Blastn. So FastViromeExplorer ran 3 times faster than ViromeScan and 16 times faster than Blastn. For the largest data set with 40 million reads, FastViromeExplorer took 2 hours and 27 minutes, a 2.5x speedup compared to ViromeScan that took 6 hours and 23 minutes. Taken together, when using NCBI virus and phage database as reference, FastViromeExplorer takes on average about 1 minute to process one million reads; when using a larger database (125,842 mVCs, 2GB), FastViromeExplorer takes 3–4 minutes to process one million reads, a 2–3x speed up compared to ViromeScan. Note that the indexing time (for both FastViromeExplorer and ViromeScan) was not counted in the running time shown (Figure 2) as indexing is only one time process. Once the index file is generated, it can be used to annotate any metagenomic data.

Simulated datasets were initially used to compare the annotation performance of FastViromeExploree with Viromescan and Blastn. Sinces viruses mutate fast, even if it is the same viral species, the viral sequences in the metagenomic data might not be exactly the same as their sequences in the reference database, it is therefore important to examine the performance of an annotation tool taking into account virus’ high mutation rate. We therefore simulated four data sets with different mutation rates (3%, 5%, 7%, and 10%) from the references and applied FastViromeExplorer, ViromeScan, and Blastn. Figure 3 shows the F1 score (recall and precision are given in supplementary file 1). All the tools have high precision (99%) across all the data sets. But as mutation rate becomes higher, the number of mapped reads reduces and recall becomes lower for all the tools. In terms of F1 score, Blastn has the best score, FastViromeExplorer has similar but slightly lower score, and ViromeScan has the lowest score. For the data set with the highest mutation rate 10%, the F1 scores for Blastn, FastViromeExplorer and ViromeScan are 0.79, 0.7, and 0.43 respectively. But FastViromeExplorer took 2 minutes compared to Blastn 8 minutes. Therefore, for these simulated data sets and using all eukarytoic viruses as the reference database, FastViromeExplorer runs four times faster than Blastn while maintaining a similar F1 score to Blastn. As viruses are known to have fast mutation rates, Blastn and its variants (e.g., Blastp) are considered the “gold standard” approach to annotate viral sequences in metagenomic data but very time-consuming, having similar performance yet running much faster is highly desired for an annotation tool.

**Figure 3:**
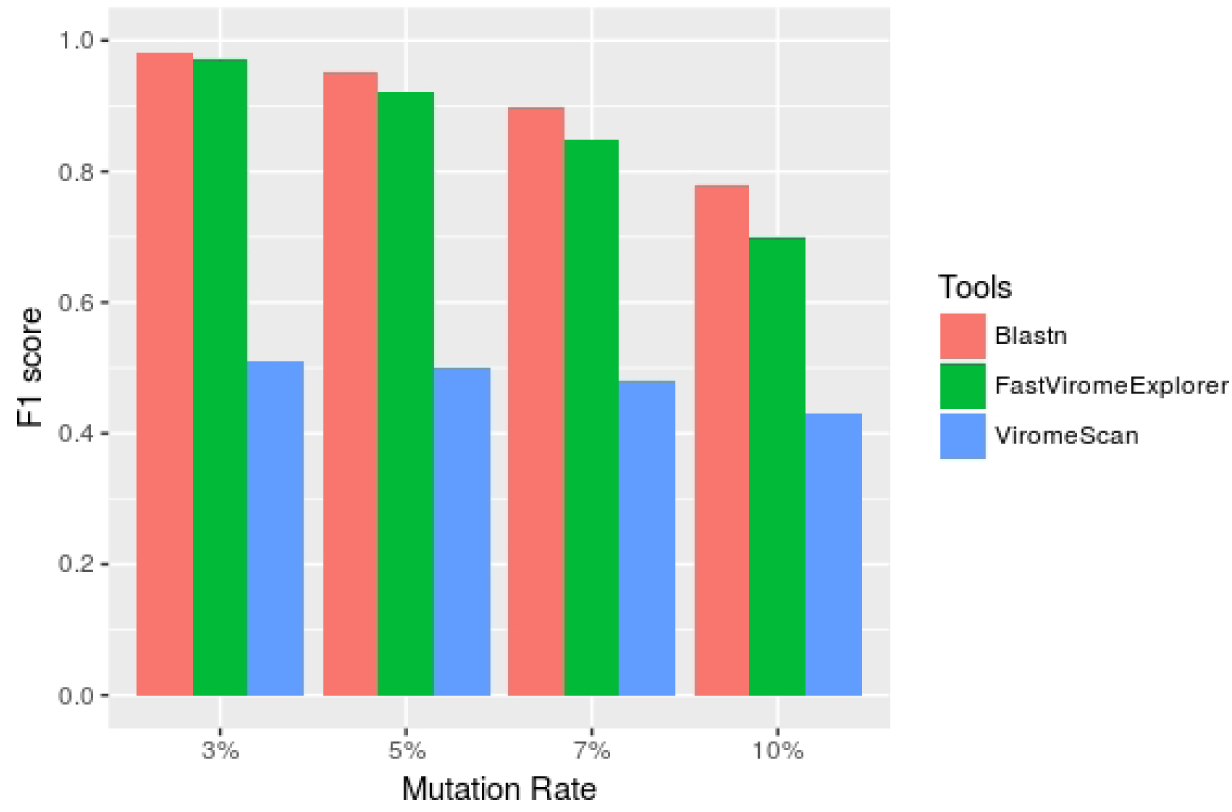
F1 score of FastViromeExplorer, ViromeScan, and Blastn when using NCBI eukaryotic viruses as the reference database and four simulated data sets of 1 million reads each with mutation rate 3%, 5%, 7%, and 10% respectively.

To examine the annotation performance of FastViromeExplorer on real data, we applied FastViromeExplorer to the fecal metagenomic samples collected from Lee et al. [14]. Lee et al. followed the dynamics and consequence of fecal micriobial transplantation (FMT) by examining the metagenomics data from a donor’s and recipients’ preFMT and postFMT samples. They constructed 92 bacterial metagenome-assembled genomes (MAGs) from reads of the donor samples and examined the occurrence of the MAGs in the recipient samples. They found that the bacterial MAGs that were present in the donor samples and also colonized the recipient samples after FMT mostly belonged to the order *Bacteroidales*. Here we examined the dynamics of viruses/phages to see whether it is consistent with the finding of Lee et al. [14].

From the annotation result of FastViromeExplorer using the 8,957 NCBI RefSeq viral genomes as reference, we observed that only three viruses (Human endogenous retrovirus K113, Glypta fumiferanae ichnovirus segment C10, and Lactococcus prophage bIL311) were found in all four donor samples, with human endogenous retrovirus K113 being the most abundant for samples 1, 3, and 4, and Lactococcus prophage bIL311 the most abundant in sample 2. For the recipient, 30 viruses were found in the preFMT sample whereas only five were found in the two postFMT samples. Among the five viruses, only Lactococcus prophage was also found in one donor sample. But as this prophage was also present in the preFMT sample, we cannot conclude that the virus was transferred from the donor to the recipient. Overall, the annotation result reveals no clear evidence of virus/phage transfer from the donor to the recipient.

We also applied ViromeScan to the fecal samples with its default reference database containing 4,370 eukaryotic DNA/RNA viruses. ViromeScan identified 847 viruses in all the samples. Compared to ViromeScan’s reference database, ours is two times bigger and it is thus surprising that ViromeScan identified a lot more viruses than FastViromeExplorer. Analysis of the ViromeScan result shows that the most abundant virus, Encephalomyocarditis virus, has all the reads mapped to a repeat region of its genome (see supplementary figure 3), indicating that the annotation is likely false positive. In fact, Encephalomyocarditis virus was also present in the initial result produced by FastViromeExplorer, but was discarded after the first filtering step. To further examine the effect of our three filtering criteria, we applied them to the ViromeScan result. Figure 4 shows that most of the viruses were filtered out and only Human endogenous retrovirus K113 and Glypta fumiferanae ichnovirus remained, both of which were also present in the final result of FastViromeExplorer. The finding here shows the importance of the filtering criteria in removing viruses that might be annotation artifacts caused by repeats, low coverage, and small genome sizes.

**Figure 4:**
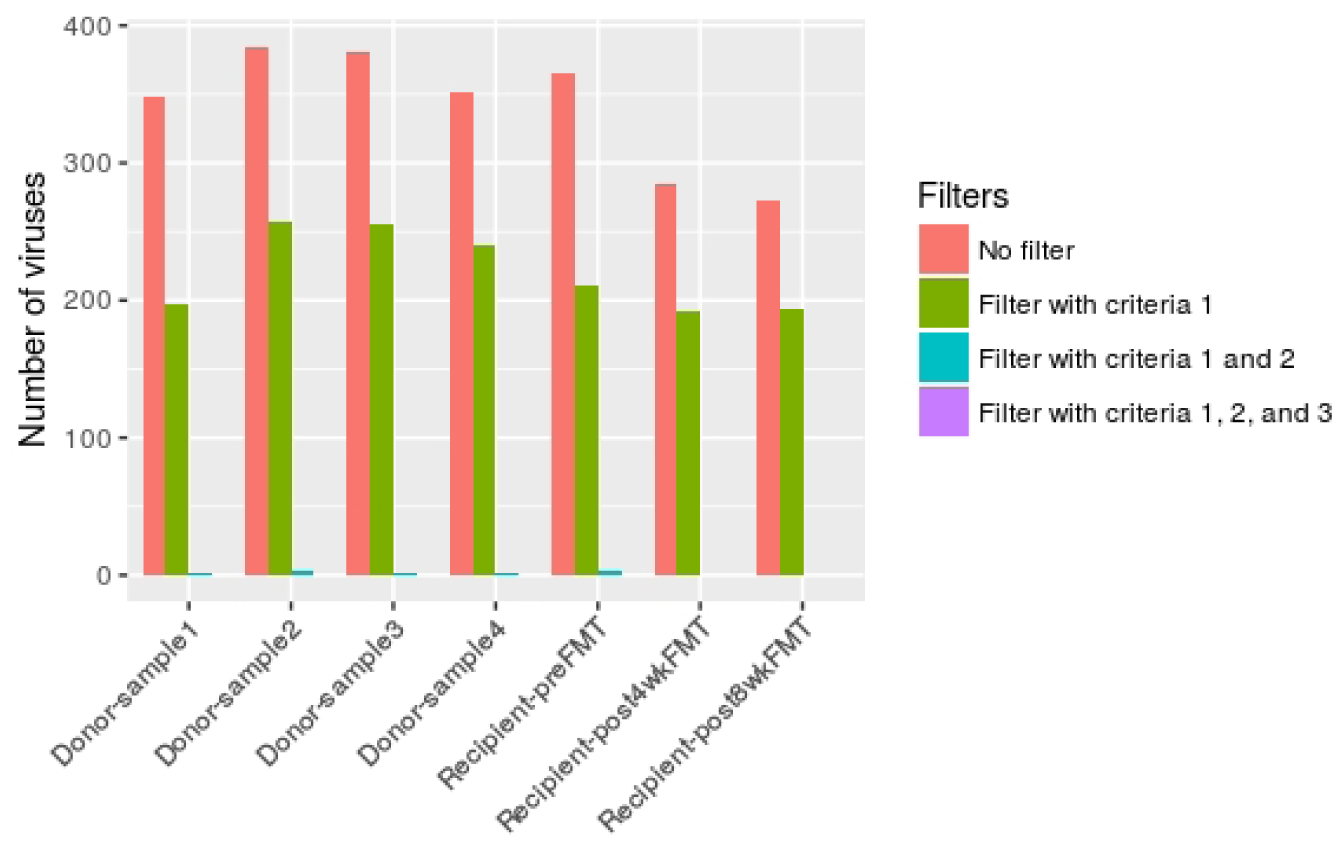
Number of viruses from ViromeScan result before applying any filter, after applying criterion 1, after applying criteria 1 and 2, and after applying all three criteria.

Since the annotation of the fecal samples using the default NCBI viral database did not reveal anything meaningful about fecal microbiota transplantation from the donor to the recipient, we tried FastViromeExplorer again using the 125,842 metagenomic viral contigs (mVCs) collected from Paez et al. [18] as reference. These mVCs are mostly unknown partial or complete viral genomes but have been predicted/annotated for their possible hosts. Therefore, the host information of the mVCs can be used to examine the annotation result. Figure 5 shows the relative abundance of host bacteria across all donor and recipient samples. The order *Bacteroidales* is more abundant than the order *Clostridiales* in all donor samples. For the recipient, prior to FMT, the order *Clostridiales* clearly dominated the microbiota, however, after the transplantation, the abundance of the order *Bacteroidales* increased dramatically and the abundance of the order *Clostridiales* decreased greatly. This result indicates that due to the transplantation, phages with host bacteria from the order *Bacteroidales* were transferred from the donor to the recipient. For example, in donor samples, “SRS049900_LANL_scaffold_14438” is one of the most abundant mVC, being the most abundant in donor samples 1 and 2, and the second most abundant in samples 3 and 4. This mVC was not present in the recipient’s preFMT sample but was highly abundant in the postFMT samples, suggesting the successful transferring of the mVC from the donor to the recipient. As the host of this mVC is from the order *Bacteroidales*, this suggests the successful colonization of bacteria from the order *Bacteroidales* from the donoor to the recipient. Therefore, our result on phage transfer following the FMT is consistent with the observation on bacterial colonization following the FMT shown in the original study [14]. The detailed annotation result is given in supplementary file 2.

**Figure 5:**
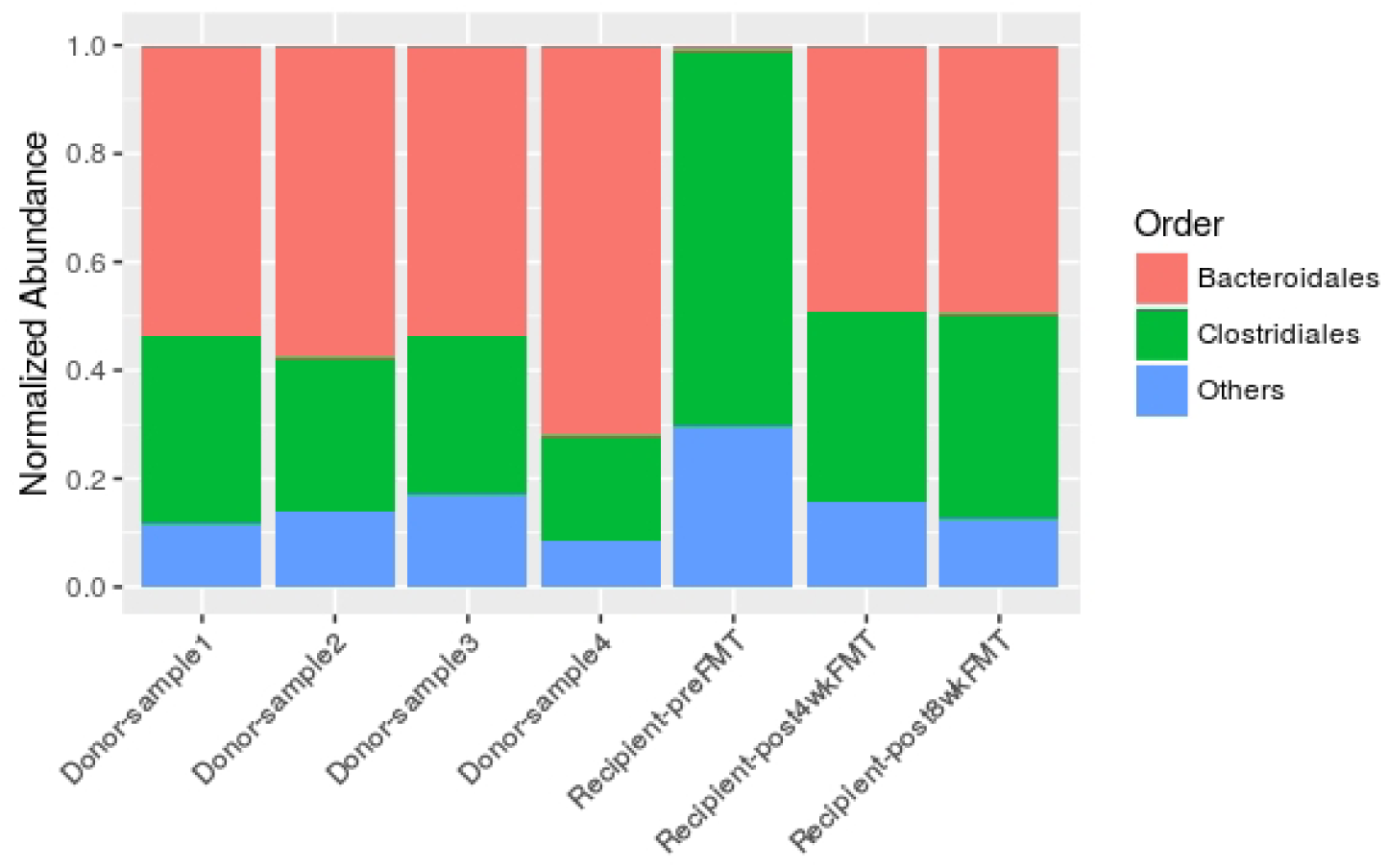
Relative abundance of host bacteria at Order level in the samples from FastViromeExplorer result using the 125,842 mVCs, where abundance is normalized by the total abundance of viruses in the sample.

Consequently, when we applied FastViromeExplorer to the samples using a larger reference database, a much clear correlation between our results and the biological results reported in the original paper emerges, which indicates the importance of of having a fast annotation and quantification tool that can easily handle bigger and more complete reference databases.

## 4 Conclusion

In this paper, we develop a new tool FastViromeExplorer for annotating virus types and abundances in metagenomic data. Worth emphasizing is that FastViromeExplorer can annotate both viruses and phages depending on the reference database users deploy. As FastViromeExplorer can process millions of reads within minutes while having similar annotation accuracy to the gold standard tool Blastn, it empowers researchers that have limitted computing power to process large metagenomic data within reasonable time. Similar to all other reference database based tools, the limitation of FastViromeExplorer is that it cannot identify a virus or phage if it is not in the reference database. Together with the result of FMT data, it highlights the pressing issue of building and/or extending the current viral sequence database for improving virus/phage annotation in metagenomic data.

## References

[1] Stephen F. Altschul, Thomas L. Madden, Alejandro A. Schäffer, Jinghui Zhang, Zheng Zhang, Webb Miller, and David J. Lipman. Gapped BLAST and PSI-BLAST: a new generation of protein database search programs. Nucleic Acids Research, 25(17):3389–3402, 1997.

[2] Nicolas L. Bray, Harold Pimentel, Páll Melsted, and Lior Pachter. Near-optimal probabilistic RNA-seq quantification. Nature Biotechnology, 34(5):525–527, 2016.

[3] Gubio S. Campos, Antonio C. Bandeira, Silvia I. Sardi, et al. Zika virus outbreak, Bahia, Brazil. Emerg Infect Dis, 21(10):1885–6, 2015.

[4] Miles W. Carroll, David A. Matthews, Julian A. Hiscox, Michael J. Elmore, Georgios Pollakis, Andrew Rambaut, Roger Hewson, Isabel García-Dorival, Joseph Akoi Bore, Raymond Koundouno, et al. Temporal and spatial analysis of the 2014-2015 Ebola virus outbreak in West Africa. Nature, 524(7563):97–101, 2015.

[5] Arthur P. Dempster, Nan M. Laird, and Donald B. Rubin. Maximum likelihood from incomplete data via the EM algorithm. Journal of the royal statistical society. Series B (methodological), pages 1–38, 1977.

[6] Sridevi Devaraj, Peera Hemarajata, and James Versalovic. The human gut microbiome and body metabolism: implications for obesity and diabetes. Clinical Chemistry, 59(4):617–628, 2013.

[7] Laura Fancello, Didier Raoult, and Christelle Desnues. Computational tools for viral metagenomics and their application in clinical research. Virology, 434(2):162–174, 2012.

[8] Jane A. Foster and Karen-Anne McVey Neufeld. Gut-brain axis: how the microbiome influences anxiety and depression. Trends in Neurosciences, 36(5):305–312, 2013.

[9] Stephen K. Gire, Augustine Goba, Kristian G. Andersen, Rachel SG Sealfon, Daniel J. Park, Lansana Kanneh, Simbirie Jalloh, Mambu Momoh, Mohamed Fullah, Gytis Dudas, et al. Genomic surveillance elucidates Ebola virus origin and transmission during the 2014 outbreak. Science, 345(6202):1369–1372, 2014.

[10] Bart L. Haagmans, Said H.S. Al Dhahiry, Chantal BEM Reusken, V. Stalin Raj, Monica Galiano, Richard Myers, Gert-Jan Godeke, Marcel Jonges, Elmoubasher Farag, Ayman Diab, et al. Middle east respiratory syndrome coronavirus in dromedary camels: an outbreak investigation. The Lancet Infectious Diseases, 14(2):140–145, 2014.

[11] Jo Handelsman, Michelle R. Rondon, Sean F. Brady, Jon Clardy, and Robert M. Goodman. Molecular biological access to the chemistry of unknown soil microbes: a new frontier for natural products. Chemistry & Biology, 5(10):R245–R249, 1998.

[12] Patrick W. Laffy, Elisha M. Wood-Charlson, Dmitrij Turaev, Karen D. Weynberg, Emmanuelle S. Botté, Madeleine JH van Oppen, Nicole S. Webster, and Thomas Rattei. HoloVir: a workflow for investigating the diversity and function of viruses in invertebrate holobionts. Frontiers in Microbiology, 7, 2016.

[13] Ben Langmead and Steven L. Salzberg. Fast gapped-read alignment with Bowtie 2. Nature Methods, 9(4):357–359, 2012.

[14] Sonny T.M. Lee, Stacy A. Kahn, Tom O. Delmont, Alon Shaiber, Özcan C. Esen, Nathaniel A. Hubert, Hilary G. Morrison, Dionysios A. Antonopoulos, David T. Rubin, and A. Murat Eren. Tracking microbial colonization in fecal microbiota transplantation experiments via genome-resolved metagenomics. Microbiome, 5(1):50, 2017.

[15] Yang Li, Hao Wang, Kai Nie, Chen Zhang, Yi Zhang, Ji Wang, Peihua Niu, and Xuejun Ma. VIP: an integrated pipeline for metagenomics of virus identification and discovery. Scientific Reports, 6, 2016.

[16] Folker Meyer, Daniel Paarmann, Mark D’Souza, Robert Olson, Elizabeth M. Glass, Michael Kubal, Tobias Paczian, A. Rodriguez, Rick Stevens, Andreas Wilke, et al. The metagenomics RAST server-a public resource for the automatic phylogenetic and functional analysis of metagenomes. BMC Bioinformatics, 9(1):386, 2008.

[17] David Paez-Espino, I. Chen, A. Min, Krishna Palaniappan, Anna Ratner, Ken Chu, Ernest Szeto, Manoj Pillay, Jinghua Huang, Victor M. Markowitz, et al. IMG/VR: a database of cultured and uncultured DNA Viruses and retroviruses. Nucleic Acids Research, 45(D1):D457–D465, 2017.

[18] David Paez-Espino, Emiley A. Eloe-Fadrosh, Georgios A. Pavlopoulos, Alex D. Thomas, Marcel Huntemann, Natalia Mikhailova, Edward Rubin, Natalia N. Ivanova, and Nikos C. Kyrpides. Uncovering Earths virome. Nature, 536(7617):425–430, 2016.

[19] Nadège Philippe, Matthieu Legendre, Gabriel Doutre, Yohann Couté, Olivier Poirot, Magali Lescot, Defne Arslan, Virginie Seltzer, Lionel Bertaux, Christophe Bruley, et al. Pandoraviruses: amoeba viruses with genomes up to 2.5 mb reaching that of parasitic eukaryotes. Science, 341(6143):281–286, 2013.

[20] Simone Rampelli, Matteo Soverini, Silvia Turroni, Sara Quercia, Elena Biagi, Patrizia Brigidi, and Marco Candela. ViromeScan: a new tool for metagenomic viral community profiling. BMC Genomics, 17(1):165, 2016.

[21] Simon Roux, Joanne B. Emerson, Emiley A. Eloe-Fadrosh, and Matthew B. Sullivan. Benchmarking viromics: an in silico evaluation of metagenome-enabled estimates of viral community composition and diversity. PeerJ, 5:e3817, 2017.

[22] Simon Roux, Francois Enault, Bonnie L. Hurwitz, and Matthew B. Sullivan. VirSorter: mining viral signal from microbial genomic data. PeerJ, 3:e985, 2015.

[23] Simon Roux, Michaël Faubladier, Antoine Mahul, Nils Paulhe, Aurélien Bernard, Didier Debroas, and François Enault. Metavir: a web server dedicated to virome analysis. Bioinformatics, 27(21):3074–3075, 2011.

[24] Simon Roux, Jeremy Tournayre, Antoine Mahul, Didier Debroas, and François Enault. Metavir 2: new tools for viral metagenome comparison and assembled virome analysis. BMC Bioinformatics, 15(1):76, 2014.

[25] Lorian Schaeffer, Harold Pimentel, Nicolas Bray, Páll Melsted, and Lior Pachter. Pseudoalignment for metagenomic read assignment. Bioinformatics, page btx106, 2017.

[26] Nicola Segata, Levi Waldron, Annalisa Ballarini, Vagheesh Narasimhan, Olivier Jousson, and Curtis Huttenhower. Metagenomic microbial community profiling using unique clade-specific marker genes. Nature Methods, 9(8):811–814, 2012.

[27] Arian FA Smit, Robert Hubley, and P. Green. RepeatMasker, 1996.

[28] Charlotte Soneson, Katarina L. Matthes, Malgorzata Nowicka, Charity W. Law, and Mark D. Robinson. Isoform prefiltering improves performance of count-based methods for analysis of differential transcript usage. Genome Biology, 17(1):12, 2016.

[29] Duy Tin Truong, Eric A. Franzosa, Timothy L. Tickle, Matthias Scholz, George Weingart, Edoardo Pasolli, Adrian Tett, Curtis Huttenhower, and Nicola Segata. MetaPhlAn2 for enhanced metagenomic taxonomic profiling. Nature Methods, 12(10):902–903, 2015.

[30] K. Eric Wommack, Jaysheel Bhavsar, Shawn W. Polson, Jing Chen, Michael Dumas, Sharath Srinivasiah, Megan Furman, Sanchita Jamindar, and Daniel J. Nasko. VIROME: a standard operating procedure for analysis of viral metagenome sequences. Standards in Genomic Sciences, 6(3):421, 2012.

[31] Laurence Zitvogel, Romain Daillère, María Paula Roberti, Bertrand Routy, and Guido Kroemer. Anticancer effects of the microbiome and its products. Nature Reviews Microbiology, 2017.

